# Including both sexes in *in vivo* research does not necessitate an increase in sample size: a key role for factorial analysis methods

**DOI:** 10.1101/2022.09.29.510061

**Authors:** Benjamin Phillips, Timo N. Haschler, Natasha A. Karp

## Abstract

In recent years, there has been a strong drive to improve the inclusion of animals of both sex during *in vivo* research, driven by a need to improve sex representation in fundamental biology and drug development. This has resulted in inclusion mandates by funding bodies and journals, alongside numerous published manuscripts highlighting the issue and providing guidance to scientists. However, progress is slow and blockers to the routine use of both sexes remain. From a statistical and experimental design perspective, concerns include difficulty selecting and conducting an appropriate analysis and the perceived need for a higher sample size to achieve an equivalent level of statistical power. When both sexes are included, analysis errors are frequent, including inappropriate pooling or sex-disaggregation of the data. These mistakes result in a failure to properly account for the variation in the data that arises from sex, and subsequently lead to poor inference regarding the biological impact of sex. The purpose of this manuscript is to address frequently cited blockers and analysis errors, thus providing a practical guide to support scientists in the design of in vivo studies which include both sexes. Primarily, we demonstrate that there is no loss of power to detect treatment effects when splitting the sample size across sexes in most common biological scenarios, providing that the data are analysed appropriately. In the rare situations where power is lost, the benefit of understanding the role of sex outweighs the power considerations. When estimating a generalisable translatable effect, where exploring sex differences are not the primary scientific objective, we recommend splitting the sample size across male and female mice as a standard strategy. We also demonstrate an optimal analysis pipeline for analysing data gathered using both sexes which is designed to help address common analysis errors.

## Introduction

There has been a bias towards using only a single sex in *in vivo* research. Though there is variation between sub-disciplines, this strategy has tended to result in a heavy bias towards male animals. For example, in 2009, only 26% of studies used both sexes and amongst the remainder there was a male bias in 80% of studies (Beery & Zucker, 2011). The negative consequences of these shortcomings on scientific enterprise are beginning to be better understood as evidence emerges that our current fundamental biological knowledge base may be biased. For example, a recent report concluded that the fundamental molecular basis of pain is highly sex dimorphic, yet much of our knowledge in this area is derived from studies solely using male organisms (Mogil, 2020). This situation risks generating a knowledge imbalance that persists through the research pipeline, ultimately manifesting in the clinic.

This issue has resulted in a push to include both male and female animals in *in vivo* studies, including mandates from funding bodies, to improve the generalisability of studies and improve the translation of studies from animals to humans. Over time, the proportion of studies including both sexes has improved (Rechlin et al., 2022; Woitowich et al., 2020), with one study estimating an increase between 2009 and 2019 from 26% to 48% from a representative sample of studies (Woitowich et al., 2020). Scientists tend to be supportive of efforts to improve sex representation in biomedical research (Woitowich & Woodruff, 2019). Unfortunately, in studies where both sexes are collected, a large proportion commit errors at the statistical analysis stage (Garcia-Sifuentes & Maney, 2021). Thus, despite an overall increase in inclusion, the proportion of studies appropriately managing sex as a biological variable (SABV; see glossary) is still low (Rechlin et al., 2022). Overall, the pace of change is slow, owing to a persistent and broad range of perceived statistical and practical blockers. These include now debunked beliefs that female animals produce more variable data (Becker et al., 2016; Beery, 2018; Prendergast et al., 2014), institution-level ingrained cultural belief about the value of studying two sexes (Karp & Reavey, 2019; Waltz et al., 2021), and a skill-gap in handling data collected under factorial designs (Garcia-Siguentes & Maney., 2021, Johnson et al., 2014). There is also a general belief that it is necessary to greatly increase the experimental sample size (N) when investigating treatment effects in two sexes (Fields, 2014; Waltz et al., 2021). Though this misconception has been addressed previously (Beery, 2018), it remains widespread and there is a need for a comprehensive exploration of the issue. Moreover, it must continue to be addressed, as at present it is a significant blocker as it drives a misguided belief that there is a trade-off between pursuing the 3Rs by means of reducing animal usage on the one hand and designing more generalisable studies on the other (Waltz et al., 2021). Thus, scientists believe that including both sexes will introduce a significant ethical, practical, and financial burden (https://www.ukri.org/wp-content/uploads/2022/03/MRC-090322-SexInExperimentalDesign-SummaryReport.pdf).

Current practices tend to strive for standardised studies by tightly controlling factors included in the experiment (e.g., using a single strain or sex of animal or including only a small age range). This reduces variability in the data but at the expense of generalisability, as the entire population of interest is under-represented in the study design. Any of a wide range of factors including animal strain, age, health status or others could be the focus of a campaign to improve research generalisability. However, sex is a particularly pressing and timely direction for focus with regards to improved representation since females account for such a large proportion of the population of interest but are currently largely overlooked. The main intent underpinning SABV strategy is to improve the generalisability of *in vivo* studies by expanding the inference space to both sexes when studying a treatment. This does not require designing experiments explicitly to test for sex differences rather estimating an average treatment effect that is generalisable to both sexes. When a large sex dimorphism is present, it is detectable when experiments are designed under this framework.

This manuscript is designed to act as a resource for scientists to implement the SABV philosophy by achieving two key goals: 1) demonstrate that studies designed to detect treatment effects including two sexes do not need to substantially increase the experimental sample size across a broad range of conditions and circumstances. 2) demonstrate an appropriate analytical pipeline including visualisation and statistical techniques for data collected from both sexes. These goals are linked insofar as appropriate statistics mitigates the need for an increased N to maintain statistical power.

When considering sex-inclusive research, the following misconceptions and data analysis errors have been reported:

- Misconception 1: designs that include both sexes will require a doubling of sample size to achieve the same power. [e.g. MRC survey, https://www.genderscilab.org/blog/three-years-in-sex-as-a-biological-variable-policy-in-practice-and-an-invitation-to-collaborate], https://www.ncbi.nlm.nih.gov/pmc/articles/PMC7957378/]
- Misconception 2: Belief that the possibility of sex effects (either a baseline differences or a treatment effect that depends on sex) will increase variability and consequently require an increased N to maintain the power (Field et al., 2014, Waltz et al., 2021).
- Error 1: Inappropriate pooling of males and females data for a treatment group to estimate an average effect (i.e. combining the data from both sexes and ignoring sex as a factor in the analysis) (Garcia-Sifuentes & Maney., 2021).
- Error 2: Disaggregation of the data by sex and independent statistical comparison between the control and treated group. Then comparing the p-values from the independent tests (Garcia-Sifuentes & Maney., 2021, Will et al., 2017).
- Error 3: Incorrect groups in statistical comparison: comparison of treated males and females (Woitowich et al, 2020).

Of these, the misconceptions (1 and 2) contain empirical claims, and for these we have constructed a range of simulated datasets that are designed to closely resemble plausible biological scenarios to assess the effect on statistical power and hence the impact on the N needed. The consequences of pooling male and female data and disaggregating the data by sex (error numbers 1 & 2) are also explored through simulation. Critically, the SABV vision does not require studies to be powered to assess whether the treatment effect depends on sex. Rather, it is sufficient to power studies to detect treatment effects of interest, whilst retaining the ability to detect large, clinically meaningful sex differences (NIH Policy on Sex as a Biological Variable., 2022, UKRI Sex in experimental design., 2022). The errors (1-3) can be addressed through guidance on appropriate analysis methods for factorial experimental designs. For this, we provide an example data analysis pipeline for processing data from studies using both sexes. A glossary of common statistical terms used within this manuscript has been developed see Box 1.

Through simulations, we have addressed the listed common misconceptions, when estimating a treatment effect that generalises to both sexes, to demonstrate that including both male and female animals does not reduce statistical power across a wide range of biologically plausible conditions. We consider the situation where power is lost to be a) rare and b) indicative of a sex dimorphism that it would be important to be aware of. Additionally, we compare statistical analysis strategies and find that inappropriate methods, including pooling the data from males and females and disaggregation of the data by sex, result in a loss of statistical power. Therefore, the importance of adopting a factorial analysis method is central to appropriately handle data from studies which include both sexes.

## Results and discussion

### Introduction to simulations to explore the impact of sex-related biological differences on statistical power

It is often believed that the inclusion of both sexes diminishes the power to detect treatment effects by introducing extra variability in the data. Thus, a study would need an increased N to detect an identical treatment effect if it included both sexes. If this were true, it would raise both ethical and cost implications for studies intending to test both sexes. We have systematically investigated this claim by the construction of simulated datasets to explore a range of possible sex-related outcomes, starting with the most biologically likely. In these simulations, we construct datasets with defined differences in the size of main effects (sex and treatment) and interactions. By repeating this construction process multiple times and testing the resultant data with a factorial analysis (Box 2), we can calculate the statistical power of each scenario by returning the proportion of times a statistically significant result occurs. In addition, we compare the power to that obtained from applying a Student’s *t*-Test where the data for each treatment has been pooled across the sexes. When a significant interaction is present it is common to conduct post-hoc pairwise statistical tests (e.g., a statistical comparison between untreated and treated animals of each sex). Therefore, we include an evaluation of the power of the post-hoc tests. Notably, the post-hoc test power taken in isolation is equivalent to the strategy of disaggregating the data by sex and conducting two independent Student’s *t*-Tests.

**Box 2:**
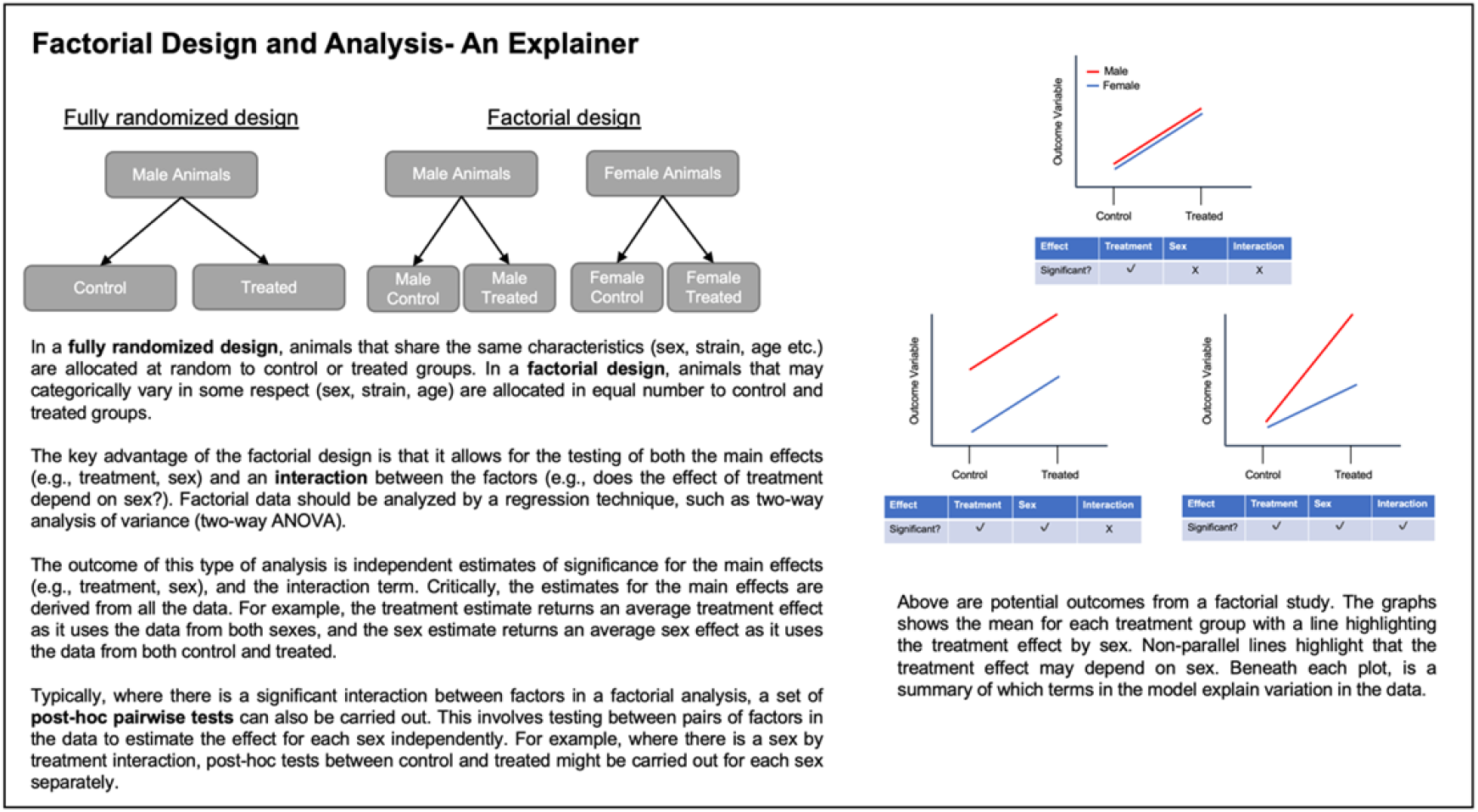
Factorial design and analysis explained.

### Scenario 1: Impact on statistical power of a baseline sex difference

In the first scenario, we tested the effect of increasing baseline sex difference (none = 0, small = 0.5, large = 1 added to the males) on the ability to detect a main effect of treatment (effect size: 0-1 added to both sexes) (Fig.1A). This is representative of the most common situation observed in studies run including both sexes (Karp et al., 2017). As the effect size of the baseline sex differences increased, there was no change in statistical power when a factorial statistical method was adopted (Fig 1B). However, there was a loss in power when the data were pooled and a Student’s *t*-Test applied (error 1)(Fig 1B). This is because factorial analysis is accounting for variability in the data that arose from a baseline difference in the sexes whereas the effect is not accounted for in the pooled analysis. This highlights the error that is committed when data from both sexes are pooled, resulting in significant loss of power to detect treatment effects.

**Figure 1:**
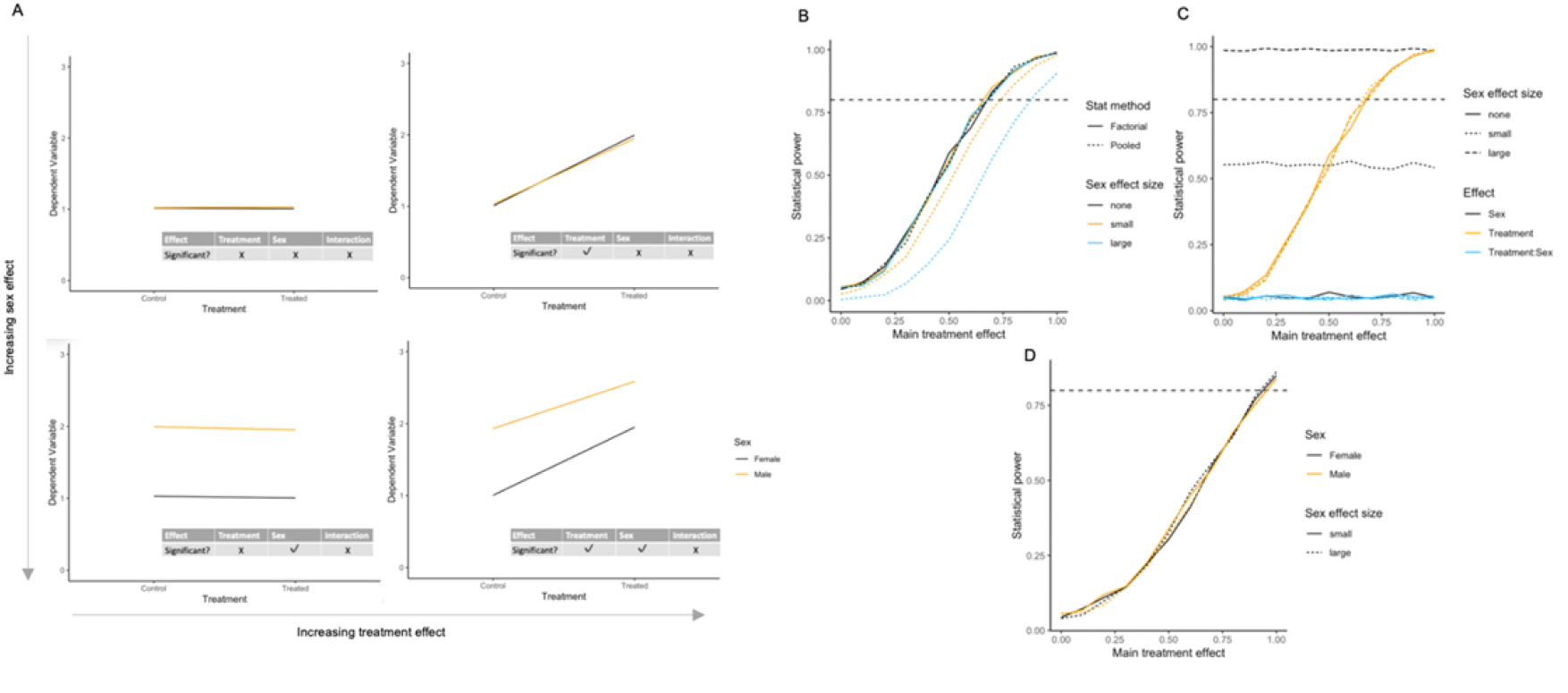
Results of simulations to explore impact of a baseline sex differences on statistical power. A) Illustrative plots of constructed datasets, ranging from zero to maximum baseline sex differences and treatment effect. B) Statistical power from both a factorial and pooled analysis to detect a main effect of treatment when the main effect of sex is either (none = 0, small = 0.5, large = 1). C) Power for each model term within factorial analysis output. D) Power for post-hoc comparison of control vs treated within each sex independently. Simulation N=1000 for each scenario assessed. For the graphs of power (B, C, D), the horizontal dashed line indicates the target power.

As the treatment effect was added to both sexes equally no biological interaction occurred. Consequently, there was no power to detect an interaction effect across the range of treatment and sex main effects. The power to detect a baseline sex difference increased as the effect size increased (Fig 1C). The power for post-hoc tests increased equivalently for both sexes as the overall treatment effect increased (Fig 1D). Notably, since the power for the post-hoc treatment test is equivalent to the power of analysing the two sexes independently, the loss of power when disaggregating the data is evident from comparison of Fig 1D and 1B. This loss of power is not of concern, because the interaction was not significant, and the treatment effect should be evaluated as a main effect. Furthermore, where there is no baseline sex effect in this scenario, the power to detect a treatment effect is identical between the factorial and pooled strategy. Therefore, there is no statistical cost to using the more complex analysis strategy when there is no variance attributable to sex since the treatment effect is estimated using all the data from both sexes. Overall, therefore, there is no loss of power to detect a treatment effect when there is a baseline sex effect in the data.

### Scenario 2: Impact on statistical power when there is a treatment effect of different sizes for the two sexes

To understand the impact of an interaction between treatment and sex on the power, where the interaction was in the same direction as a main effect of treatment, datasets were constructed to allow an exploration of the impact of an increasing interaction effect (none = 0, small = 0.5, large = 1) in the presence of a main effect of treatment (varied from 0 to 1)(Fig. 2A). There was no baseline sex difference in the data. When the treatment by sex interaction increased (Fig 2A), the power to detect a main effect of treatment increased in line with the size of the interaction effect (Fig 2B). As the interaction size increased, the pooled analysis (error 1) lost power to detect the main effect of treatment compared to the factorial analysis, as there was an increase in sex-related variance not accounted for. This highlights that the false belief that sex-related variance reduces statistical power to detect a treatment effect results from inappropriate analysis. There was an increase in power to detect a treatment by sex interaction as the interaction effect size increased (Fig 2C). There was greater power to detect a post-hoc effect in the sex which exhibited the largest treatment effect, which increased as a function of effect size (Fig 2D).

**Figure 2:**
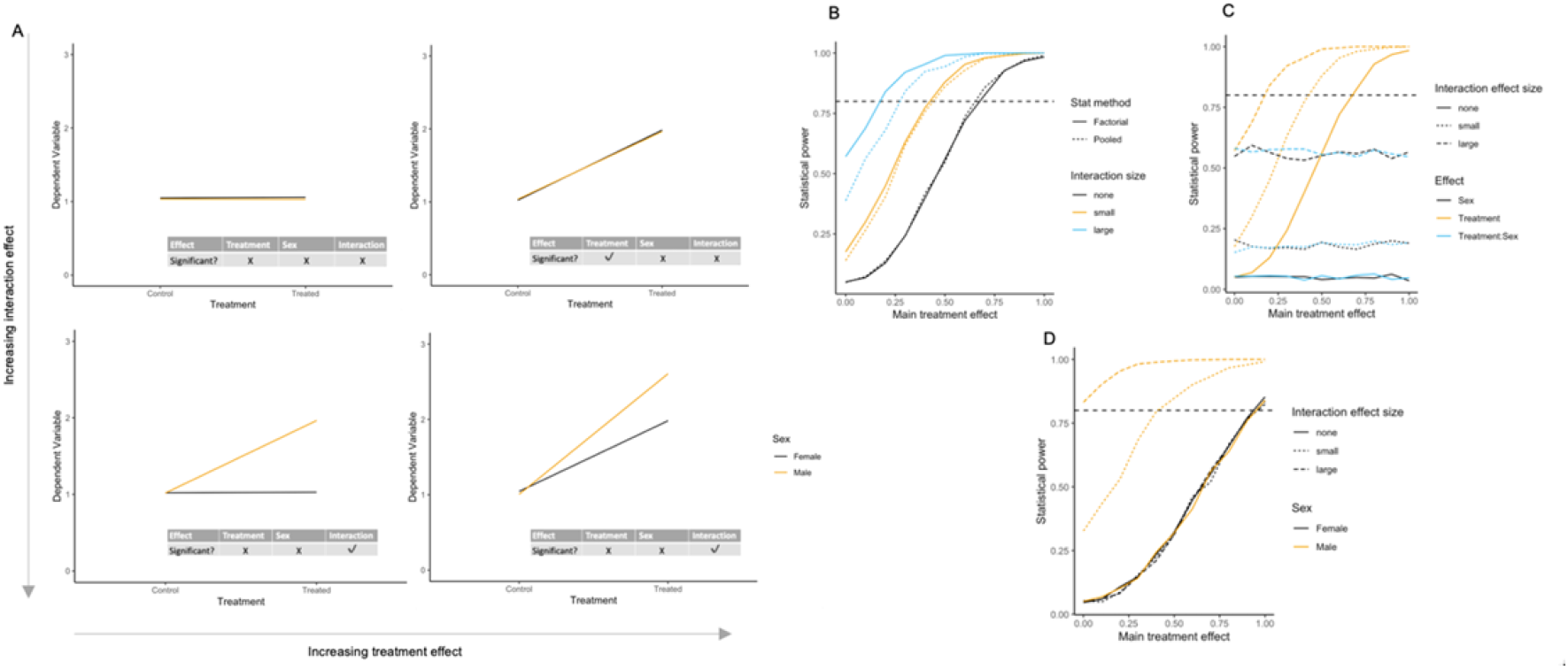
Results of simulations to calculate statistical power where there is an interaction between treatment and sex in the data where the interaction effect is in the same direction as the main treatment effect. A) Illustrative plots of constructed datasets, ranging from no effect to maximum treatment by sex interaction (none = 0, small = 0.5, large = 1) for varying sizes of a main treatment effect (0-1). B) Statistical power to detect treatment effect per size of interaction effect and statistical method. C) Power for each model term within factorial analysis output. D) Power for post-hoc comparison control vs treated within each sex. Simulation N=1000 for each scenario assessed. For the graphs of power (B, C, D), the horizontal dashed line indicates the target power.

These simulations highlight the error of disaggregating the data and conducting independent Student’s *t-*Tests on each sex separately. The factorial workflow estimates the main effects and tests the interaction. If the interaction effect is biologically and statistically significant, then the post-hoc testing allows an assessment of treatment effect independently for each sex. Conversely, if pairwise tests are carried out without an initial factorial analysis (error 2: disaggregation of data by sex) there is no ability to assess for a treatment by sex interaction and therefore is a statistical error as it precludes the ability to support a claim of a differential treatment effect. This is because it is a statistical error to compare two independent statistical tests (e.g. a significant and non-significant result) and claim a difference on this basis (Makin et al., 2019, Nieuwenhuis et al., 2011). Thus, a factorial analysis is required to properly implement the SABV philosophy and to avoid misinterpretation of the data.

### Scenario 3: Impact on statistical power when there is a treatment effect in one sex only

An extreme example of a treatment by sex interaction is when there is a treatment effect in one sex only (Fig 3A). In this scenario, there was a power benefit to using a factorial analysis compared to the pooling strategy as the interaction effect size increased (Fig 3B). This is because the increased between-sex variance was not accounted for when the data were pooled. In this scenario, the power to detect the main effects of sex and treatment and the treatment by sex interaction were identical as the interaction effect size increased (Fig 3C). There was only power to detect a post-hoc effect in the sex which exhibited the treatment effect, which increased as the interaction effect size increased (Fig 3D). Thus, there is a power benefit when a factorial analysis is adopted compared to pooling (error 1). However, in the case where the sex exhibiting the treatment effect would have been studied in isolation and the entire N used within a single sex, there was a loss of power to detect a treatment effect when splitting the N across the two sexes (Fig 3E). In this rare scenario (Karp et al., 2017), the conclusions obtained by studying only a single sex would have been skewed. If only the sex showing the large treatment effect was included in the study, the researcher may have mistakenly assumed that the conclusion generalised to the entire population. Conversely, if only the sex displaying low sensitivity to treatment were included, we may erroneously conclude that the treatment was ineffective. Therefore, in the rare scenario in which power to detect an overall main effect of treatment is reduced by splitting the intended N across the two sexes, our knowledge of the treatment effect is increased and unbiased when using factorial analysis.

**Figure 3:**
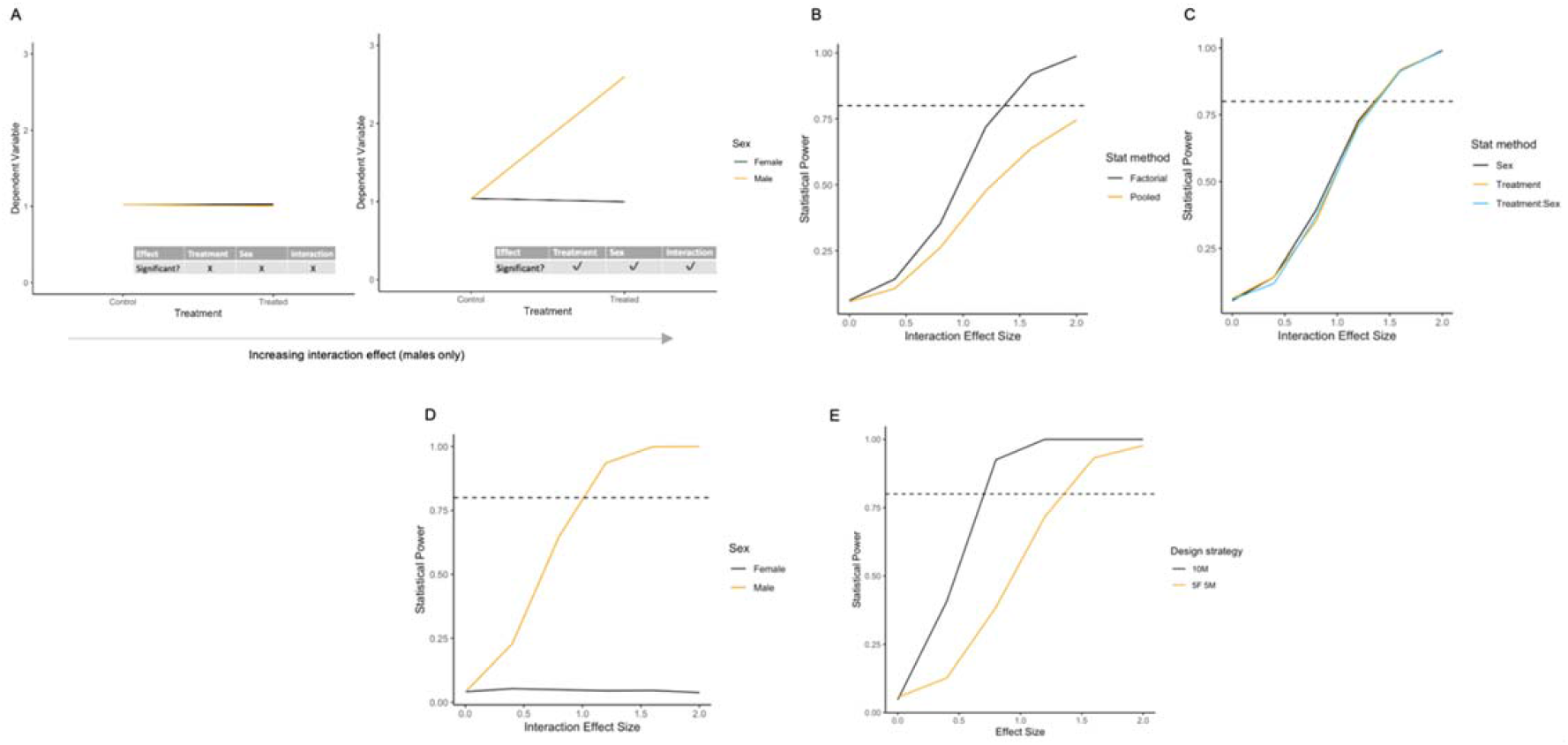
Results of simulations to calculate statistical power where there is an interaction driven by a treatment effect in one sex only. A) Illustrative plots of constructed datasets, ranging from a zero to maximum treatment by sex interaction effect (0-2). B) Statistical power to detect the main effect of treatment effect as a function of the interaction effect size for each statistical method. C) Power for each model term within factorial analysis output. D) Power for post-hoc comparison control vs treated within each sex. E) Power for contrasting design strategies-10 males vs 5 females and 5 males. Simulation N=1000 for each scenario assessed. For the graphs of power (B, C, D, E), the horizontal dashed line indicates the target power.

### Scenario 4: Impact on statistical power when there is an interaction driven by opposite sex effects

Though rare biologically (Karp et al., 2017), it is possible a treatment may produce an opposite effect in each sex (Fig 4A). In this scenario, the power to detect a main effect of treatment was low as expected (Fig 4B), as the mean treatment effect across sexes was equivalent to zero. The factorial analysis produced the expected false call rate (proportion of the time that a significant result is detected when there is no true difference) for this significance threshold of approximately 5% for all effect sizes, whilst the pooling strategy false positive rate decreased as the size of the effect increased. The conservative behaviour of the pooling strategy arises from the unpartitioned variance that is attributable to sex. With a factorial analysis, there was equivalent power to detect a main effect of sex and the treatment by sex interaction, but no power to detect a main effect of treatment as the mean treatment effect averaged across sexes is approximately zero (Fig 4C). There was also equivalent post-hoc testing power to detect a treatment effect in each sex (Fig 4D). In comparison to a single sex study where the entire N was focused on one sex, the power of the factorial design when the N is split across the two sexes is lower. However, the inclusion of only a single sex in this scenario would have resulted in a highly biased conclusion that may be inappropriately generalised to the broader population of interest. A strong opposite effect would merit follow-up studies to understand the basis of the differential treatment effect.

**Figure 4:**
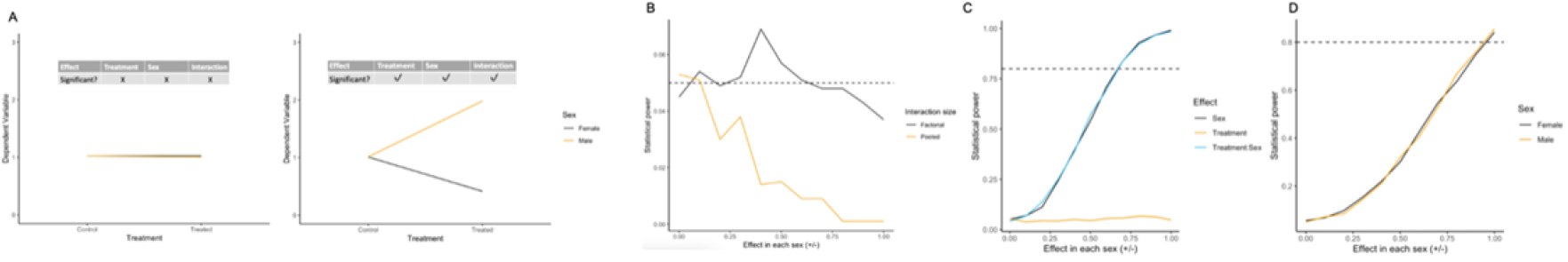
Results of simulations to calculate statistical power where there is an interaction driven by opposite treatment effect in each sex. A) Illustrative plots of constructed datasets, ranging from no effect to maximum opposite effect in each sex (0-1 male, 0 to −1 female). B) Statistical power to detect the main effect of treatment as a function of the interaction effect size for each statistical method. C) Power for each model term within factorial analysis output. D) Power for post-hoc comparison control vs treated within each sex. Simulation N=1000 for each scenario assessed. For the graphs of power (B, C, D), the horizontal dashed line indicates the target power.

### A pipeline for analysis of data collected from two sexes: a case study

The murine nephrotoxic nephritis model (NTN) recapitulates multiple features of glomerulonephritis, a form of chronic kidney disease, and is induced via administration of antibodies targeted against the murine glomerular basement membrane (GBM) (referred to as nephrotoxic serum (NTS)) (Nagai et al., 1985; Xie et al., 2004). This model may display a differential induction effect by sex, with female mice reported to display greater susceptibility to NTS (Ougaard et al., 2018). Therefore, we selected this model as a pilot to investigate an optimised analysis pipeline for analysis of data including two sexes. Notably in the clinic, anti-GBM disease (Goodpasture Syndrome) has historically been described as male-dominated, however, this has been disputed in more recent investigations, where no sex bias was shown (Beckwith et al., 2022).

#### Step 1: visualisation

As sex should be considered a primary factor of experimental interest, data should be visualised by sex separately for each treatment level. Best practice is to present individual datapoints from each animal, with indications of central tendency and variance for each treatment group (e.g., boxplot). Here we present the Urinary Albumin-to-Creatinine (UACR) (Fig 5A) alongside genetic expression data obtained from mice treated with either control (serum) or NTS, separated by sex (Fig 5B).

**Figure 5:**
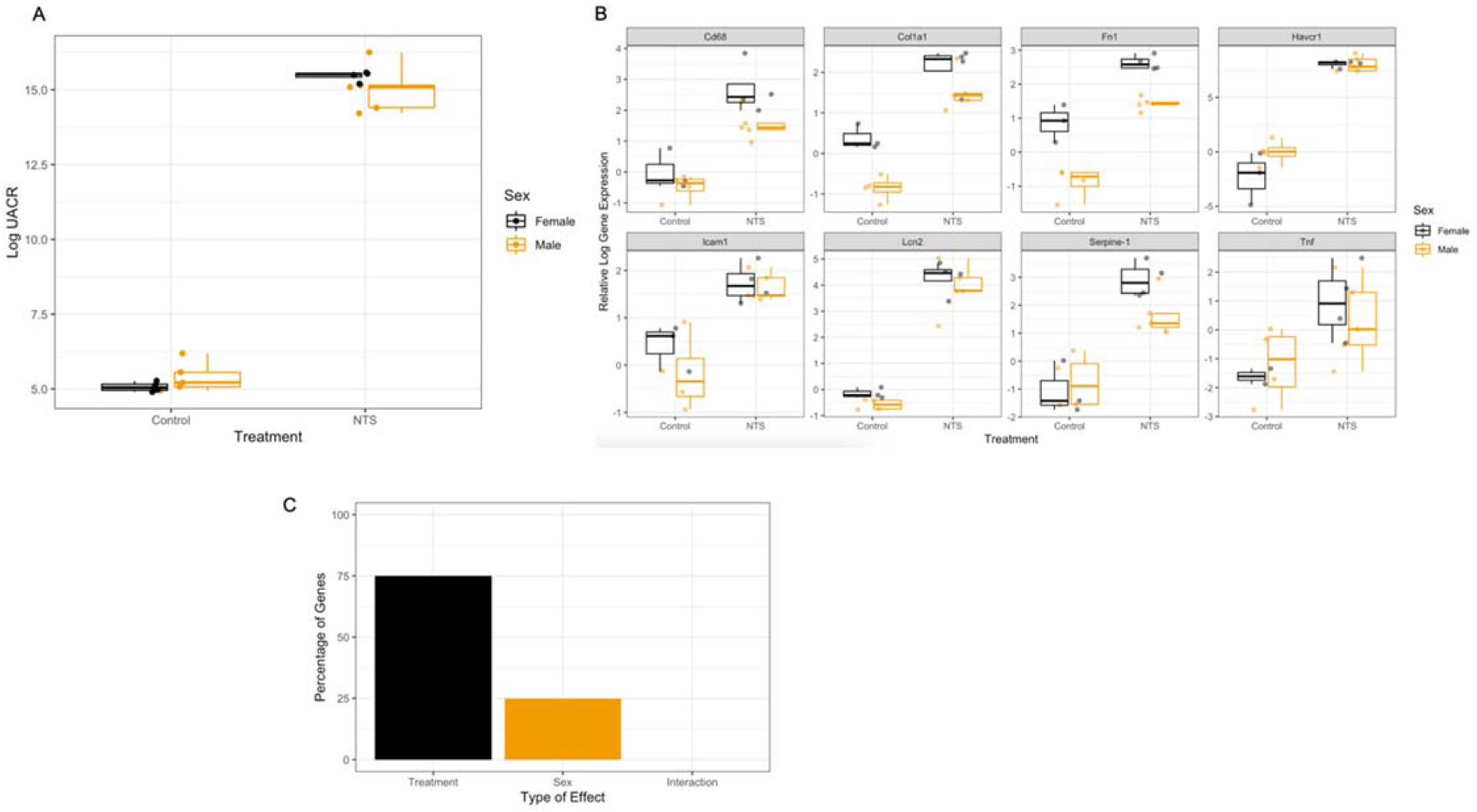
*Visualisation of data by treatment and sex in the NTN model of kidney disease. A) Urinary Albumin-to-Creatinine Ratio (UACR) at day 4 post-dosing (N = 5 per sex per treatment). B) Relative expression levels for all measures genes (N; Control female = 3, control male = 4, NTS female = 4, NTS male = 5, apart from Tnf, where control female = 2). C) Overall summary of statistically significant model effects (Benjamini-Hochberg corrected p*-value < 0.05*) in the dataset*.

#### Step 2: two-way ANOVA

Two-way ANOVA is a regression analysis technique that handles factorial experimental designs by partitioning the variance that is attributable to each factor in the data independently prior to estimating the interaction effect. This is a factorial design, as there are two factors of interest: the treatment and sex. In this example, the treatment has two levels (NTS and control, and sex has two levels (male and female). Note, this analysis method is not restricted to two levels for a factor; the treatment could have been a treatment with four levels (vehicle, low, medium, and high dose).

For UACR, there was a significant effect of NTS treatment (F(1, 14) = 1685.979, *p* < 0.0001). There was no significant main effect of sex (F(1, 14) = 0.045, *p =* 0.835) or treatment by sex interaction (F(1, 14) = 2.451, *p* = 0.14)(Fig. 5A). Therefore, we did not detect any significant sex-related effects on the primary measure of model induction.

For the panel of genes, each gene was analysed independently by two-way ANOVA. An example 2-way ANOVA output is displayed in Table 1. These results highlight that both genes exhibit a significant main effect of treatment. For *Col1a1*, there is also a significant main effect of sex.

**Table 1:** Example two-way ANOVA output when testing the three model terms (treatment, sex and interaction). Shown are genes Col1a1, which exhibited both a significant main effects of treatment and sex, and Lcn2, which exhibited a significant main treatment effect following NTS administration. Neither gene exhibited a significant treatment by sex interaction.

| **Col1a1** | Degrees of freedom | Sum of squares | Mean squares | F-value | P-value |
| --- | --- | --- | --- | --- | --- |
| Treatment | 1 | 17.629 | 17.629 | 93.961 | 0.0000005 |
| Sex | 1 | 2.986 | 2.986 | 15.917 | 0.00179 |
| Treatment * Sex | 1 | 0.409 | 0.409 | 2.178 | 0.16577 |
| Residuals | 12 | 2.251 | 0.188 |  |  |
| **Lcn2:** | Degrees of freedom | Sum of squares | Mean squares | F-value | P-value |
| Treatment | 1 | 77.88 | 77.88 | 185.195 | 0.000000011 |
| Sex | 1 | 0.72 | 0.72 | 1.723 | 0.214 |
| Treatment * Sex | 1 | 0.00 | 0.00 | 0.00 | 0.995 |
| Residuals | 12 | 5.05 | 0.42 |  |  |

Alongside statistical significance, it is important to evaluate effect sizes to interpret the results in their biological context. A common method for this is to examine contrasts through changes in estimated marginal means. An example output from this method of analysis is provided in Table 2.

**Table 2:** Example estimated marginal means analysis for treatment terms. Estimated means are presented alongside the estimated change, with standard errors and CL (confidence limits).

| **Col1a1:** | **Treatment** | Estimated marginal mean | SE | df | Lower CL | Upper CL |
| --- | --- | --- | --- | --- | --- | --- |
|  | Control | -0.238 | 0.165 | 12 | -0.598 | 0.123 |
|  | NTS | 1.822 | 0.145 | 12 | 1.506 | 2.139 |
|  | **Contrast** | Estimated change | SE | df | T ratio | P-value |
|  | Control-NTS | -2.06 | 0.22 | 12 | -9.358 | <0.0001 |
| **Lcn2:** | **Treatment** | Estimated marginal mean | SE | df | Lower CL | Upper CL |
|  | Control | -0.365 | 0.248 | 12 | -0.904 | 0.175 |
|  | NTS | 4.075 | 0.218 | 12 | 3.602 | 4.549 |
|  | **Contrast** | Estimated change | SE | df | T ratio | P-value |
|  | Control-NTS | -4.44 | 0.33 | 12 | -13.472 | <0.0001 |

The estimated changes suggest large, generalisable mean shifts for both genes following treatment (averaging across sexes). This is consistent with the metrics of significance observed from the ANOVA output, but the size of the changes, alongside estimates of variability, provide additional biological context which assists with interpretation.

Since multiple independent statistical tests are being conducted, there is the potential for accumulation of false positives and hence adjustment for multiple testing needs to be applied. Multiple methods exist (Chen et al., 2017), with varying approaches to manage the risk. The method selected should be based on the research hypothesis in question (see https://multipletesting.com/analysis for a freely accessible tool). In this example, we have implemented the Benjamini-Hochberg procedure, as it balances robust control of the false discovery rate with adequate maintenance of statistical power (Benjamini & Hochberg, 1995). The statistically significant results are displayed in Table 2.

**Table 2:**
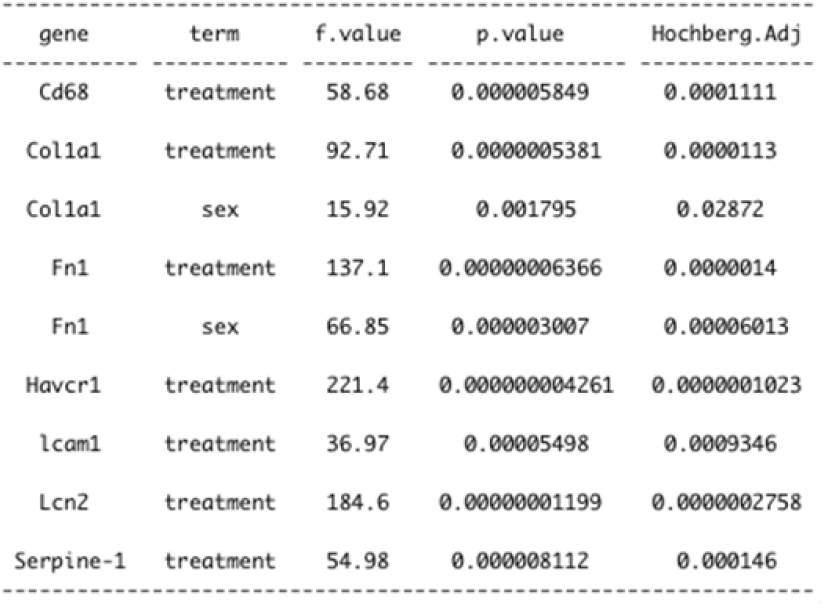
Significant results from two-way ANOVA on the gene panel (N= 8 genes). For each gene, the degrees of freedom were DF(1, 12). The BH-adjusted p-values are presented alongside the original p-values.

A summary of the significant model effects in the dataset is highlighted in Fig 5C. Several genes exhibited a significant main effect of treatment. Notably, no significant treatment by sex interaction effects were detected. A minority of genes demonstrated a baseline sex difference. We expect that this will be a typical result in many studies. The study was not powered to detect an interaction between treatment and sex, rather to give an indication of any large sex divergences in treatment effect, that may merit further investigation, were present. This is in line with the current SABV philosophy and guidance. The study has estimated the average treatment effect across the two sexes, thus is more generalisable as we have explored a wider inference space.

#### Step 3: post-hoc tests

The choice of whether to conduct post-hoc tests depends on the biology in question. For example, where a model of disease is being induced, it could be appropriate to estimate the treatment effect from the ANOVA alone, as a generalisable estimated effect. Conversely, it could be important to precisely understand the magnitude of the treatment effect in both sexes individually. For example, when there is a known difference in the biology between sexes, it would then be appropriate to estimate the effect in each sex independently using post-hoc tests.

In the present dataset no treatment by sex interactions were detected. However, where a two-way ANOVA analysis yields a significant interaction, it is typical to additionally carry out pairwise post-hoc tests of significance to estimate the treatment effect within each sex and compare the size and direction of these effect to explore the biological difference that is responsible for the statistical interaction. A frequently committed error at this stage where two sexes are included is to misidentify the appropriate pairwise comparisons (error 3). For example, researchers often erroneously compare the means of treated males to treated females which conflates baseline sex differences with differences resulting from treatment. The risk from this error is high, as baseline sex differences are common (Karp et al., 2017). Another consequence of this error is the inflation of the false positive multiplicity burden by carrying out irrelevant comparisons (e.g., treated males to untreated females). Although there may be situations where a scientific hypothesis merits other comparisons we recommend the post hoc tests compare untreated to treated groups within each sex by default. Importantly, where the number of treatment levels increases, post-hoc comparisons should also generally undergo adjustment for multiple testing, otherwise the risk of producing spurious false significance is greatly inflated.

## Conclusions

The SABV philosophy is to include both sexes to detect a major difference in response to treatment across the two sexes. This approach allows us to statistically test for treatment by sex interactions and visualise the data by both sexes. This is driven by a desire to increase the translational confidence of the results and doesn’t require experiments to be powered to detect treatment by sex effects. Our simulations show that for most biologically expected situations (where treatment effects are equivalent across the sexes or there is a baseline sex difference) there is not a loss of power. When there is a small difference in the size of the treatment by sex interaction, estimating the average effect is ideal for translational understanding as this is a generalisable conclusion. The approach does, however, lead to a loss of power when there is a large treatment by sex interaction (e.g., opposite or the effect only occurs in one sex). In this uncommon scenario and where the N has been split across the two sexes, we may fail to detect a significant treatment effect (due to the lower power) but would gain important knowledge suggestive of a large sex dimorphism. Taken together, these outcomes support the recommendation to split the intended N across the two sexes.

To facilitate progress in sex inclusion in *in vivo* research, it is crucial to provide scientists with both evidence and practical resources that challenge the blockers that currently stand in the way of change. This manuscript provides a resource intended to address the common misconception that studies including both sexes require a doubling of the sample size. To achieve this, we have performed extensive statistical simulations to evaluate power under a range of common biological scenarios when splitting the N across two sexes. Critically, we did not identify any common scenarios that result in a loss of power to detect treatment effects. Rarely, large interactions in the data may produce an appreciable decrease in treatment effect power. In these scenarios, the knowledge gained that the treatment has a differential impact between sexes outweighs the statistical consequences. If a disease affects both sexes but an effect is observed in only one, this may bring into question the validity of the model or the generalisability of the treatment. The simulations also demonstrate the pitfalls of some frequent analysis mistakes, including the inappropriate pooling and disaggregation of data collected from two sexes. Our simulation investigations have focused on the power to detect treatment effects specifically, as the SABV philosophy does not require powering to detect sex differences. The approaches above heavily depend upon the appropriate application of the correct factorial analysis methods, and it is therefore critical that laboratory scientists receive focused support in developing their statistical capabilities.

## Box 1: Glossary

Common statistical terms used within this manuscript, adapted from (Percie du Sert et al., 2020) and placed in the context of *in vivo* research are detailed below:

### Effect size

Quantitative measure of differences between groups, or strength of relationships between variables.

### Factor

Factors are independent categorical variables that the experimenters control during an experiment in order to determine their effect on the outcome variable. Example factors include sex or treatment.

### Factorial design

An experimental design that is used to study two or more factors, each with multiple discrete possible values or levels.

### Independent variable

A variable that either the experimenter controls (e.g., treatment given or time) or is a property of the sample (sex) or a technical feature (e.g., batch or cage) that can potentially affect the outcome variable.

### Interaction effect

When the effect of one independent variable (factor) depends on the level of another. For example, the observed treatment effect depends on the sex of the animals.

### Levels

Are the values that the factor can take. For example, for the factor sex the levels are male and female.

### Main effect

A main effect is the overall effect of one independent variable on the outcome variable averaging across the levels of the other independent variable.

### Outcome variable

A variable captured during a study to assess the effects of a treatment. Also known as dependent variable or response variable.

### Power

For a predefined, biologically meaningful effect size, the probability that the statistical test will detect the effect if it exists (i.e., the null hypothesis is rejected correctly). Can also be called sensitivity.

### Treatment

a process or action that is the focus of the experiment. For example, a drug treatment or a genetic modification.

### Sex as a biological variable (SABV)

the research philosophy that emphasises the importance of including both sexes in *in vivo* studies in such a way that a generalisable treatment effect is detectable. Critically, sex should be treated as a variable of primary biological interest. There is no requirement to power to detect a baseline difference between the sexes or treatment by sex interaction, but studies will detect large differences where they exist.

## Methods

### Statistical simulations

To explore the impact of sex either as a main effect or when it interacts with the treatment on power, simulation studies were run. In these studies, 1000 datasets were constructed to represent an experiment with both sexes and a control and treatment group for each scenario of interest. The resulting datasets were analysed using the statistical test of interest and statistical power was assessed, as the proportion of times a statistically significant effect was called for the element of interest, at a significance threshold p<0.05. The following characteristics were fixed: the experiment consisted of 5 animals per treatment group per sex with data sampled from a normal distribution with a baseline mean of 1 and variance of 0.5. The impact of having a main effect of treatment, a main effect of sex and/or an interaction between sex and treatment on the statistical power was then explored (Table 3). Datasets were tested with either a factorial pipeline (regression analysis in R equivalent to a two-way ANOVA) or with a Student’s *t*-Test after pooling the data across the sexes for a treatment. In addition, uncorrected post-hoc pairwise tests were applied between untreated and treated data in each sex using the R package *emmeans.* The objective of these simulations is to understand general trends and relationships as such the exact values are not relevant but represent typical animal experiments.

**Table 3:** Details of how the simulation datasets were constructed to represent various biological situations.

| Simulation | Objective | Main effect of sex | Main effect of treatment | Interaction between the treatment and sex |
| --- | --- | --- | --- | --- |
| 1 | Impact of a baseline sex difference on statistical power | Varied<br>Signal added to means of males: none (0), small (0.5) or large (1) | Varied<br>Signal added to the treatment group mean:<br>0-1 increments of 0.1 | None |
| 2 | Impact of an interaction between sex and treatment on statistical power | None | Varied<br>Signal added to the treatment group mean<br>0-1 increments of 0.1 | Varied<br>Signal added to the mean of the treated males<br>None (0), small (0.5) or large (1) |
| 3 | There is a treatment effect in one sex only | None | None | Varied<br>Signal added to males only: 0-2 in increments of 0.2 |
| 4 | There is an interaction driven by opposite sex effects | None | None | Varied<br>Signal added to both sexes symmetrically in opposite direction<br>Male: 0-1 increments of 0.1<br>Female: 0- -1 increments of 0.1 |

### *In vivo* experiments

#### General statement and licencing

All animal experiments were conducted in accordance with the United Kingdom Animal (Scientific Procedures) Act 1986 and associated guidelines, approved by institutional ethical review committees (Alderley Park Animal Welfare and Ethical Review Board; Babraham Institute Animal Welfare and Ethical Review Board) and conducted under the authority of the Home Office Project Licences (PF344F0A0). All animal facilities have been approved by the United Kingdom Home Office Licensing Authority and meet all current regulations and standards of the United Kingdom. The manuscript was prepared in accordance with the ARRIVE guidelines.

#### Animals, housing and husbandry

*C57BL6N* mice (10 male and 10 female; age 6 weeks; Taconic France) were housed in groups of five mice per cage with an inverse 12 hours day-night cycle in a controlled temperature (19-23 °C) and humidity (45±65%) room as directed in the Code of Practice for the Housing and Care of Animals Bred, Supplied or Used for Scientific Purposes. Animals were housed in individually ventilated cages (IVC) and received food (Special Diet Services R&M No 1) and water ad libitum. Within the cage, the bedding material composed Aspen Eco-pure wood chips, nesting of sizzle nest and hemp happi-mats; as environmental enrichment cardboard smart home, cardboard or red polycarbonate tunnel, and Aspen chew stick. Animal health was checked twice daily from the day of delivery and water changed twice weekly.

### Procedures

All procedures were carried out in a laminar flow hood. Male mice were always handled first when doing any procedures and hands cleaned with ethanol after handling each cage group to remove any traces of scent/odour from the previous cage to avoid any male aggression/fighting occurring.

#### Chip implantation

After a one-week acclimatization period, an identification chip was implanted by subcutaneous injection to identify the individual animals.

#### Urine collection

Urine was collected on day 14. For this, mice were placed in empty cages for a period of maximal 30min. Each mouse had assigned one cage for urine collection which was used for the whole duration of the study. After collection, urine was transferred to collection plates and stored at −80C until analysed.

#### Nephrotoxic nephritis

Five male and five female mice were either injected with vehicle (control serum; Probetex; PTX) or nephrotoxic serum (Probetex; PTX001S-MS). For this, sera were diluted 1:1 with phosphate-buffered saline and injected intravenously into the tail vein on two consecutive days (day 0 and day 1) at a dose of 5ml/kg bodyweight.

Animals were randomly allocated to a treatment group by use of an in-house minimisation randomisation tool (Altman & Bland, 2005) to ensure balanced distribution of body weight across the treatment groups. During the randomisation step, the staff were masked to the treatment information. During intervention, the support staff were aware of the treatment they were applying. During the conduct of the experiment the cages were not labelled with the treatment information and a common welfare management plan was implemented for all treatment groups. During the outcome assessment the samples were labelled with an identification number blinding masking both the treatment and group information. During the analysis data was labelled with treatment group. All data was included in analysis. The analysis followed the analysis plan predefined in the experiment planning stage.

#### Sample size justification

In-house historic data was used to estimate the expected variance for a range of experimental parameters of interest. The final sample size was selected as it balanced the ability to detect treatment effects with practical considerations across the range of parameters.

### Sample analyses

#### UACR analysis

To determine the UACR, urine was thawed on ice and subsequently, centrifuged 3min at 4000 × g at 4 °C. Urinary albumin was quantified by using a mouse albumin ELISA kit (ALPCO) according to the manufacturer's instructions. Urinary creatinine was measured by using a creatinine assay kit (Abcam) according to the manufacturer's instructions. Albumin was normalized to creatine levels and represented as mg/g.

#### RNA isolation, cDNA synthèses and qRT-PCR analyses

Total RNA was extracted with a MagMAX™-96 Total RNA Isolation Kit (Cat #AM1830) and a KingsFisher 96 Flex system (Thermo Fisher Scientific) according to the manufacturer's instructions. For this, snap-frozen kidneys were homogenized and lysed in MagMAX™-96 lysis buffer using a TissueLyser II machine (Qiagen). RNA quality and concentration were determined by a NanoDrop-1000 spectrophotometer (Thermo Fisher Scientific). One microgram of total renal RNA was transcribed to cDNA using the High Capacity RNA-to-cDNA kit (Applied Biosystems). Quantitative analysis of target mRNA expression determined by TaqMan method using the Fast Advanced Master Mix (TaqMan) and was calculated using the ΔΔCT method. TaqMan duplex analyses were conducted using an QuantStudio™ 12K Flex Real-Time PCR system (Life Technologies). Gene expression levels were normalized to glyceraldehyde-3-phosphate dehydrogenase (*Gapdh*). TaqMan probe IDs are provided below in Table 4. Due to a technical issue with qPCR machine, data from 4 animals was not obtained.

**Table 4 :** TaqMan probes.

| **Gene** | **TaqMan probe IDs** | **Dyes** |
| --- | --- | --- |
| *Gapdh* | Mm99999915_g1 | VIC |
| *Cd68* | Mm03047343_m1 | FAM |
| *Col1a1* | Mm00801666_g1 | FAM |
| *Fn1* | Mm01256744_m1 | FAM |
| *Havcr1* | Mm00506686_m1 | FAM |
| *Icam1* | Mm00516023_m1 | FAM |
| *Lcn2* | Mm01324470_m1 | FAM |
| *Serpine-1* | Mm00435858_m1 | FAM |
| *Tnf* | Mm00443258_m1 | FAM |

### Statistical analysis of *in vivo* data

The observed UACR data measured on day 4 post-dosing were log2 transformed prior to analysis by two-way ANOVA. Gene expression data obtained by qPCR were normalised to a housekeeping gene (*Gapdh*) to derive the double delta cycle threshold values (DDCT). Subsequently data were log2 transformed, and statistical analysis was applied. There were no exclusion criteria and no datapoints were removed prior to the analysis. Data were analysed by two-way ANOVA, with treatment and sex as factors. All *p*-values derived from the analysis of the qPCR data were pooled and then corrected for multiple comparison by the Benjamini-Hochberg method (Benjamini & Hochberg., 1995) using the R package “multtest”. All analysis was carried out using R version 4.1.1 in Rstudio (www.r-project.org).

## Abbreviations

SABV: Sex as a biological variable
3Rs: Replacement, reduction, refinement
N: Total sample size number
ANOVA: Analysis of variance
UACR: Urinary albumin-to-creatinine ratio
NTS: Nephrotoxic serum
NTN: Nephrotoxic nephritis

## Contribution

BP: Conceptualization, Formal analysis, Writing – Original draft, Writing – Review & Editing

TH: Data Curation, Writing – Review & Editing

NAK: Conceptualization, Project administration, Formal analysis, Writing – original draft, Writing – review & editing

## Acknowledgements

We are grateful to Lorraine Miller and Sara Dearman for technical assistance with the experimental work and Chris Heath and Esther Pearl for helpful comments on earlier drafts of the manuscript.

